# ACEseq – allele specific copy number estimation from whole genome sequencing

**DOI:** 10.1101/210807

**Authors:** Kortine Kleinheinz, Isabell Bludau, Daniel Hübschmann, Michael Heinold, Philip Kensche, Zuguang Gu, Cristina López, Michael Hummel, Wolfram Klapper, Peter Möller, Inga Vater, Rabea Wagener, ICGC MMML-Seq project, Benedikt Brors, Reiner Siebert, Roland Eils, Matthias Schlesner

## Abstract

ACEseq is a computational tool for allele-specific copy number estimation in tumor genomes based on whole genome sequencing. In contrast to other tools it features GC-bias correction, unique replication timing-bias correction and integration of structural variant (SV) breakpoints for improved genome segmentation. ACEseq clearly outperforms widely used state-of-the art methods, provides a fully automated estimation of tumor cell content and ploidy, and additionally computes homologous recombination deficiency scores.

---

Copy number aberrations (CNAs) play an important role in tumorigenesis and are often used to subgroup cancer entities. Whole genome sequencing (WGS) identifies CNAs at unprecedented resolution, but poses challenges to CNA calling algorithms such as non-random errors and coverage biases^1^. Changing degrees of genomic complexity, tumor heterogeneity, varying tumor cell content (TCC) and aneuploidy are further challenges when analyzing tumor genomes.

Many modern tools combine tumor/control coverage ratios with B-allele frequencies (BAF) of heterozygous SNPs^2,3^. Some tools correct for GC bias, a major source of noise in the coverage signal^4^, and allow for the incorporation of SV breakpoints to assist segmentation^5^. However, to the best of our knowledge, none of the available tools provides all of the above-mentioned features.

Here, we present ACEseq, a tool to estimate absolute allele-specific copy numbers on WGS data. ACEseq involves coverage bias correction, genome segmentation allowing the incorporation of previously known breakpoints, TCC and ploidy estimation, and absolute allele-specific copy number calculation to enable fully automated CNA calling on cancer WGS data without prior information requirements.

The first step of ACEseq performs coverage bias correction, which significantly reduces noise levels (Figure 1). Noisy coverage profiles as depicted in Figure 1A cause over-segmentation and can mask CNAs. While GC bias correction greatly reduces noise, a remaining fluctuation of the signal is still observed in the shown sample (Figure 1B). This fluctuation can be attributed to replication timing coverage bias, which is particularly prominent in fast-replicating tumors^6^ (Supplementary Figure 1). Due to cells in S-phase these samples show a higher average coverage in early replicating regions than late replicating regions, as the fraction of cells with already replicated DNA at early loci is higher. Correction for replication timing bias further smoothens the coverage profile considerably, enabling more robust genome segmentation in the next step (Figure 1C).

**Figure 1:**
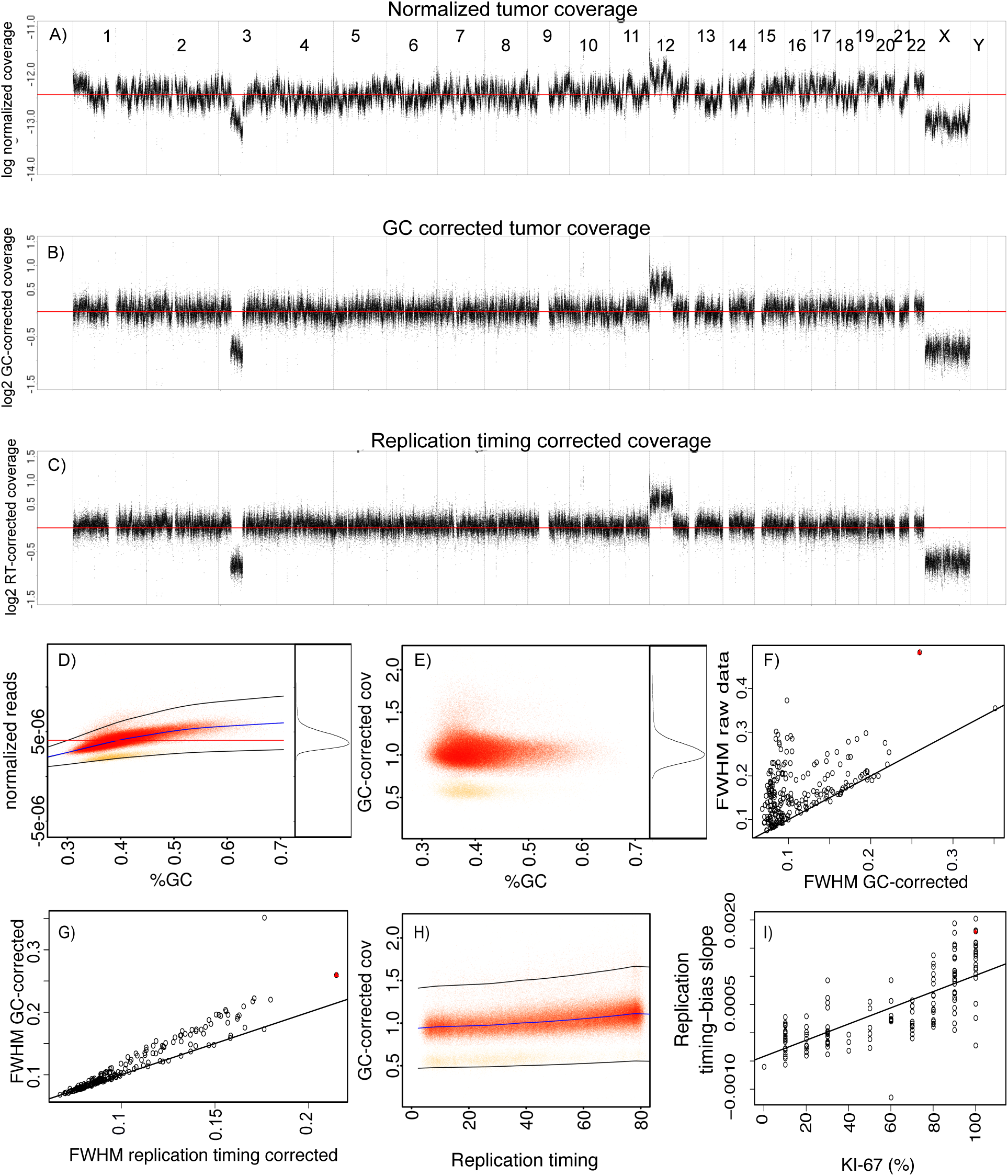
GC- and replication-timing bias correction and QC. Coverage profile for 10 kb windows before correction (A), after GC-bias correction (B), and after GC- and replication timing-bias correction (C). D: GC-bias correction curve fitted to the raw data shown in A, to firstly identify the main copy number state windows (red points) before a second curve is fitted used for correction of the bias. E: GC-corrected coverage distribution used for FWHM estimation. Comparison of FWHM estimates prior to and after GC bias (F) and replication-timing bias (G) correction for 219 lymphoma samples. H: Replication-timing bias correction curve estimating the replication speed. I: Comparison of KI-67 and estimated replication-timing slope. Sample 4112512 shown in panel A-E & H and is marked by a red triangle in F, G & I. RT: replication timing; Cov: coverage; FWHM: full width half maximum.

During the coverage bias correction steps ACEseq records statistical parameters that carry technical and biological information about the sequenced samples. Both bias correction steps are based on loess curve fitting (Supplementary Methods). Slope and curvature of the fitted GC curve (Figure 1D) indicate the magnitude of the bias. Differences in GC bias between a tumor sample and its matched healthy control indicated by these quality metrics likely affect sensitivity and specificity of the variant calling procedures for mutation types like insertions and deletions (INDELs) and single nucleotide variants (SNVs) due to differences in coverage^1^. The full width half maximum (FWHM) captures the evenness of coverage (Figure 1E, Supplementary Methods). For the vast majority of analyzed samples the FWHM decreases drastically with GC bias correction (Figure 1F, ø_reduction_=28%) and even further with additional replication-timing correction (Figure 1G, ø_reduction_=6%). The FWHM after bias corrections indicates remaining coverage fluctuations and hence serves as direct quality parameter for CNA calling. Notably, it also helps to assess the quality of sequencing libraries, a feature that has been used routinely by the International Cancer Genome Consortium PanCancer Analysis of Whole Genomes (ICGC PCAWG) community^7^. Experimental proof for the reliability of the slope from the loess curve fitted for replication-timing correction (Figure 1H) as estimator of the tumor proliferation rate could be demonstrated with KI-67 estimates, where we could show a significant correlation (n=147 germinal center derived B-cell lymphomas, p-value < 0.01, Figure 1I).

We often observed extremely noisy coverage profiles in matched controls from projects outside the ICGC MMML-Seq, possibly due to wrong handling of blood samples, preventing accurate copy number calls based on tumor/control ratios. For such samples ACEseq offers an option to replace the coverage signal from the matched control with an independent control whilst still maintaining the BAFs of the matched control. This control replacement option enables full analysis of these sample pairs including reliable discrimination between runs of homozygosity (ROH) in the germline and somatic loss of heterozygosity (LOH). Furthermore ACEseq can be run without matched control enlarging the spectrum of samples that can be processed.

Parallel to coverage correction SNPs are haplotype-phased to increase the sensitivity of allelic imbalance detection. Subsequently, ACEseq segments the genome based on changes in the BAF and tumor/control coverage ratio. Previously known breakpoints from SV calling algorithms such as DELLY^8^ or SOPHIA (manuscript in preparation) can be incorporated. Resulting raw genomic segments are clustered and merged when indicated to reduce oversegmentation (Supplementary Figure 2, Supplementary Methods). Final segments are used for TCC, ploidy and copy number estimation. The final copy number data are visualized for inspection and validation of the analysis (Supplementary Figure 4). Additionally, genomic measures well known to be significantly associated with homologous recombination (HR) defects are computed: the homologous recombination deficiency (HRD)-, the large scale transition (LST)^9^- and the telomeric allelic imbalances (TAI)^10^-score. A connection between these parameters and treatment response to platinum containing neoadjuvants and poly-ADP-ribose polymerase (PARP) inhibition has been recently indicated^11–14^.

**Figure 2:**
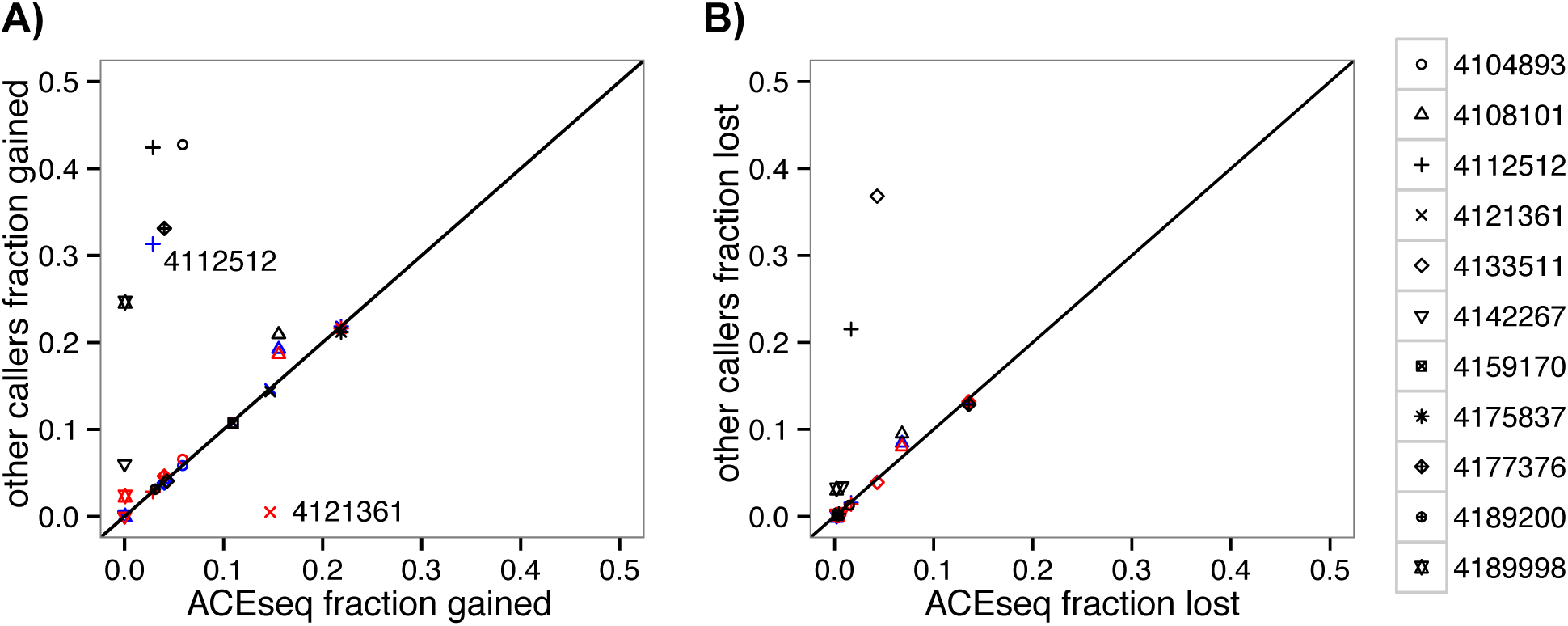
Comparison of gained (A) and lost (B) fraction of the genome for 11 B-cell lymphoma samples. The fraction called by ACEseq is compared to the other three callers. Two samples that deviate strongly for ABSOLUTE and TITAN are marked with the sample ID. TITAN: blue, THetA: black, ABSOLUTE: red.

To evaluate ACEseq’s performance we compared it to ABSOLUTE^3^ (SNP array) as well as TITAN^15^ and THetA^16^ (WGS) using 11 B-cell lymphoma samples from the ICGC MMML-seq project^17^ selected based on the availability of SNP array data from germline DNA. First, we compared ploidy and TCC predictions of the tools (Supplementary Table 1). Fluorescence *in situ* hybridization (FISH) analyses were taken as gold standard for ploidy estimations. Here, ACEseq, TITAN and ABSOLUTE showed very similar concordance with FISH based ploidy assessments. Since no gold standard for TCC was available, a comparison with an orthogonal method was used: the median mutant allele frequencies (MAF) from somatic SNVs in diploid balanced regions (Supplementary Methods). While this measure can be affected by the presence of subclonal SNVs, it can be considered as a lower boundary for the true TCC. We observed that ACEseq was able to predict the TCC with highest accuracy compared to the other tools based on the number of samples deviating less than 10% from MAF-based estimates (Supplementary Table 1).

Next we compared fractions of the genome with copy number gain and loss (Figure 2). Most tools reported similar fractions of gains and losses with the exception of THetA. THetA deviated strongly from the other methods in several samples, probably due to strong differences in TCC estimations. TITAN and ABSOLUTE only deviated from the ACEseq results in one sample each. A further investigation of these revealed that sample 4121361 was estimated at much higher TCC by ABSOLUTE, which requires a larger change in coverage for a segment to be called as gain. The other sample (4112512), called with higher fraction of gains by TITAN, was strongly affected by replication timing bias. Though ploidy and TCC were estimated at similar levels, the concordance of allelic as well as total copy number level was very low (Supplementary Table 2). TITAN and THetA showed a considerably higher number of segments than ACEseq (factor 2-5x higher) for this particular sample, suggesting that replication timing-dependent coverage bias led to oversegmentation. Resulting small segments, reflecting peaks and valleys of the noisy raw coverage track, would then be assigned to different copy number states. Titan and THetA increased the fraction of amplified genome from less than 3 % as determined by ACEseq and ABSOLUTE to 29% and 42%, respectively. No indications for this reported increased fraction were found in the raw coverage data (Figure 1A). Furthermore, the sample’s karyotype (46,X,- X,del(3)(p14p24),t(8;14)(q24;q32),+del(12)(q15)[10]/46,XX[1]), matched the ACEseq copy number predictions, demonstrating better estimates by ACEseq. Only ABSOLUTE showed results similar to ACEseq for this sample, though its TCC estimation was 68% below the ACEseq and the MAF-based estimate.

For a more detailed comparison we calculated the overall concordance of copy number calls for both total and allele-specific copy numbers (Supplementary Table 2). Strikingly, the highest concordance was observed between ACEseq and ABSOLUTE (average agreement Ø=0.96) emphasizing the robustness of the copy number calls as these methods use a different, independent data basis with WGS and SNP arrays, respectively. The concordance of ACEseq with the WGS-based methods was much smaller (average agreement: THetA Ø=0.45, TITAN Ø=0.84), though it increases to 0.91 for TITAN upon removal of the fast replicating sample 4112512 further confirming ACEseq copy number calls. Results from THetA differed substantially from ACEseq in 7 out of 11 samples, in which THetA reported much higher fractions of the genome as gained or lost. Again this is probably related to the strongly deviating TCC estimates and over-segmentation.

Overall, these results demonstrate the good performance of ACEseq in fully automated TCC and ploidy as well as allele-specific copy number estimation. ACEseq clearly benefits from its unprecedented integration of many and partially new features into a single tool. Even though ABSOLUTE and TITAN performed on a similar level for many of the samples they bear several shortcomings. ABSOLUTE always offered multiple TCC/ploidy solutions, reaching up to more than 40 possible solutions for one sample. The desired solution had to be extracted manually from an R-object and required further manual interaction. TITAN resulted in very good TCC estimation with the downside that the ploidy needs to be set in advance. Testing different ploidies requires multiple runs per sample. Additionally a strong replication timing bias caused problems for TITAN leading to over-segmentation and subsequently larger fractions of segments assigned as a gain or loss.

In conclusion, ACEseq provides a novel analysis platform for fully automated CNA calling on cancer WGS data without the requirement of prior information or any necessity for manual interference. By integrating GC and replication timing bias correction it improves segmentation and CNA calling performance compared to other tools. Importantly, it further provides quantitative metrics, which have been widely used for automatized quality control in large-scale pan cancer WGS projects. ACEseq is comprehensively documented under aceseq.readthedocs.io and freely available at https://github.com/eilslabs/ACEseqWorkflow.

## Acknowledgement

We thank P. Ginsbach and R. Drews for their contribution to establish impute2 and ABSOLUTE analyses, L. Fischer for improving the tool efficiency and Thomas Wolf for statistical support. This work was supported by the BMBF-funded Heidelberg Center for Human Bioinformatics (HD-HuB) within the German Network for Bioinformatics Infrastructure (de.NBI) (#031A537A, #031A537C), and the BMBF-funded projects ICGC MMML-Seq (#01KU1002A-J) and ICGC DE-MINING (#01KU1505E,G). Infrastructural support of the KinderKrebsInitiative Buchholz/Holm-Seppensen zu R.S. is gratefully acknowledged.

## Author contributions

K.K, I.B, D.H, Z.G and M.S developed and implemented ACEseq. Integration into the Roddy pipeline framework was done by K.K, M.H. and P.K. The ICGC MMML Seq network coordinated by R.S. provided sequencing, FISH, SNP-array and pathology data. In particular, M.Hu., W.K., P.M., provided KI-67 measurements; C.L. performed FISH analyses and karyotyping, I.V. and C.L. provided SNP array data, R.W. coordinated quality control of sequencing data. K.K. analyzed data. R.E., R.S., B.B. and M.S. supervised the research. K.K. and M.S. wrote the manuscript. All authors provided feedback on the manuscript.

